# Revealing Transcriptomic Responses in *Escherichia coli* During Early Antibiotic Exposure

**DOI:** 10.1101/2025.09.17.676618

**Authors:** Yuan Yuan, Ying Hefner, Richard Szubin, Jaemin Sung, Bernhard O. Palsson

## Abstract

The earliest responses of pathogenic bacteria to antibiotics can affect the outcome of an infection. While long-term adaptations have been extensively studied, the immediate transcriptional changes that unfold immediately following antibiotic exposure remain poorly understood. Here, we applied iModulon analysis to time-resolved transcriptomic data from *Escherichia coli* exposed to subinhibitory concentrations of two antibiotics (ampicillin and ciprofloxacin), capturing transcriptional regulatory changes occurring within the first 30 minutes of exposure. This analysis reveals an integrated, three-phase response: an immediate and sustained primary response that broadly activates stress programs, a transient secondary response that restores redox balance, and a tertiary response that supports long-term survival through metabolic remodeling and antibiotic-specific defenses. These results highlight a coordinated and dynamic regulatory strategy describing how metabolic, redox, and stress responses are integrated to manage the physiological challenges of antibiotic stress. By disentangling these overlapping transcriptional regulatory programs, this work offers a genome-scale understanding of how survival mechanisms unfold during the critical moments following antibiotic exposure. The study opens new directions for investigating early survival mechanisms and the possible identification of new targets to disrupt the initial adaptation process.

**Importance:** Initial bacterial responses to antibiotics are important for survival and can influence the development of tolerance and resistance. Yet this period remains poorly understood, in part because the transcriptional responses that unfold within minutes of antibiotic exposure are complex and difficult to interpret. In this study, we applied novel data generation and data analytics approaches to untangle the complexity of the initial response of *Escherichia coliI* to two antibiotics. We reveal a three-phase process that explains how *E. coli* coordinates stress responses, maintains redox homeostasis, and establishes longer-term defenses. The novel transcriptomic analytics elucidate independently regulated sets of genes that constitute cellular processes. By identifying all such cellular processes that react over the initial time scale, we can deconvolute the response based on first principles of cellular physiology.

## Introduction

The rise of antimicrobial resistance presents an urgent global health crisis, driving the need for deeper insights into how bacteria sense, respond, and adapt to antibiotic exposure^1^. While decades of research have revealed key aspects of bacterial defense mechanisms, much of our understanding comes from observations made hours after exposure—long after the earliest cellular decisions have shaped the outcome. These early moments of antibiotic exposure may hold key insights into how bacteria initiate their defensive responses and transition toward longer-term adaptations, potentially revealing new strategies for therapeutic intervention.

Bacterial responses to antibiotics involve complex, coordinated changes in gene expression across multiple cellular systems^2–5^. Traditional transcriptomic analyses have identified diverse stress responses, metabolic adjustments, and resistance mechanisms, but they often struggle to disentangle the transcriptional regulatory networks (TRNs) that orchestrate these changes. Advances in high-quality RNA sequencing and computational methods have enabled systematic approaches to uncover regulatory structure from gene expression data. Among these advancements, iModulon analysis has emerged as a powerful tool for understanding microbial TRNs^6–12^. iModulons are independently modulated gene sets, derived from the Independent Component Analysis (ICA) of transcriptomic data. They capture the activity of specific regulatory modules within the cell^13,14^. By tracking iModulon activities across conditions, we can observe coordinated shifts in gene expression that reflect specific transcriptional responses^15,16^. This framework is particularly powerful for understanding how multiple regulatory systems operate simultaneously and dynamically during stress responses, making it well-suited for dissecting the complex cellular response to antibiotics.

In this study, we employ iModulon analysis to investigate the transcriptional landscape of *Escherichia coli* K-12 strain MG1655 during the first 30 minutes of exposure to sub-inhibitory concentrations of two antibiotics. By capturing gene expression from as early as 1.5 minutes post-exposure, we reveal a rapid, highly coordinated response that unfolds in three overlapping, yet distinct, phases. The primary response is characterized by an immediate and sustained activation of broad stress-related programs, reflecting a generalized effort to stabilize cellular function under sudden antibiotic pressure. As this heightened activity strains the cell’s redox balance, a transient secondary response is triggered to restore homeostasis through temporary activation of anaerobic pathways. Alongside these adjustments, a tertiary response starts to guide longer-term adaptations, marked by targeted remodeling of metabolic regulation and the emergence of antibiotic-specific defenses, together helping the cell optimize survival under continued stress. These phases operate on overlapping but different timescales and serve distinct purposes, and together form a cohesive framework that enables *E. coli* to manage the immediate impact of antibiotics while transitioning toward sustained adaptation. This three-phase process outlined by iModulons offers new insight into how bacterial regulatory networks orchestrate early responses to antibiotic exposure, capturing the dynamic interplay of stress mitigation and long-term survival strategies.

## Results

### iModulons Capture Gene Expression Variations Caused by Antibiotic Exposure

We investigated the transcriptional response of *E. coli* MG1655 to two antibiotics: ampicillin and ciprofloxacin. Ampicillin, a β-lactam antibiotic, inhibits cell wall synthesis, leading to cell lysis, while ciprofloxacin, a fluoroquinolone, targets DNA gyrase and topoisomerase IV, disrupting DNA replication and repair. Gene expression was profiled at 0, 1.5, 3.5, 7.5, 15, and 30-minutes post-exposure. For ampicillin, we examined two concentrations (1/4x MIC and 1/16x MIC) to assess dose-dependent transcriptomic effects. After quality control, 35 antibiotic-treated samples were included in the analysis. To enhance the robustness of our machine learning-based iModulon analysis, these 35 samples were combined with the MG1655 samples from the PRECISE 1K dataset^15^, and ICA was applied to the combined set of 613 RNA-Seq samples (Figure 1a, b).

**Figure 1.**
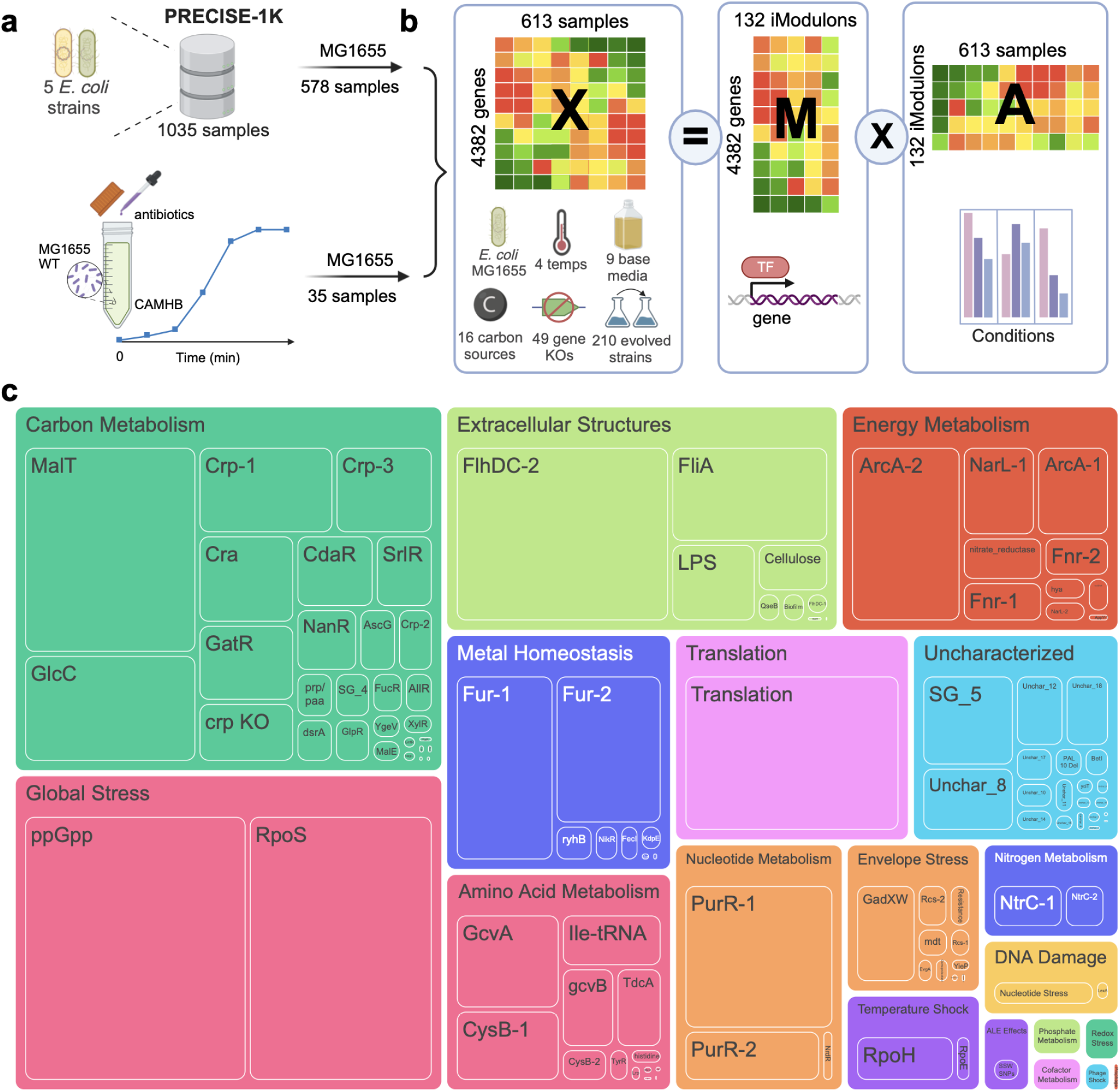
Overview of Dataset and iModulon Results. **a)**. Dataset composition. The dataset used in this study combines 578 MG1655 samples from the PRECISE-1K dataset and an additional 35 samples from the antibiotics exposure time series generated in this study. **b)**. ICA decomposes the gene expression matrix **X** into two separate matrices: the iModulon matrix **M** and the activity matrix **A**. The columns of the **M** matrix contain the independent components, from which the gene membership of iModulons is identified. The rows in the **A** matrix represents the activity of each iModulon across the different conditions in the dataset. Parts of panels a). and b). are created with Biorender.com^17^ **c)**. Treemap showing all the iModulons and their assigned functional categories. The size of the boxes represent the variance the iModulon explains in gene expression in the antibiotic exposure samples. iModulons that are identified as artifacts from data combination and normalization are removed from this figure (See Supplementary Note 1).

This analysis yielded 132 iModulons, accounting for 84% of global expression variation and 86.9% of the variation in the antibiotic study. Antibiotic exposure triggers diverse regulatory responses represented by iModulons across multiple functional categories. The explained variance of each iModulon can be calculated to quantify its contribution to expression variation in our dataset. A treemap of all the iModulons is shown in Figure 1c where the box size indicates the explained variance of the iModulon in the antibiotic exposure samples. Stress response iModulons contribute substantially to the expression variation in antibiotic-exposed samples, followed by iModulons associated with various metabolic processes, motility and biofilm formation, and energy metabolism. The full iModulon table with iModulon categories and explained variance can be found in Supplementary Table 1.

### The Response to Antibiotics is Global and Multi-layered

Detailed analysis of iModulon activities reveals that *E. coli*’s early response to antibiotics is rapid, global, and multi-layered. The majority of iModulons exhibit activity changes at different time points following exposure, reflecting a dynamic and phased response. To characterize this process, we categorized iModulons with similar activity patterns and identified a three-phase structure of the early response (Figure 2a). The initial transcriptional changes are largely non-specific and occur in two rapid phases: a primary response, which represents an immediate reaction to stress, and a secondary response, which emerges as a redox consequence of the primary response. The tertiary response, in contrast, reflects a transition toward long-term transcriptomic remodeling, including antibiotic-specific regulatory changes (Figure 2b).

**Figure 2.**
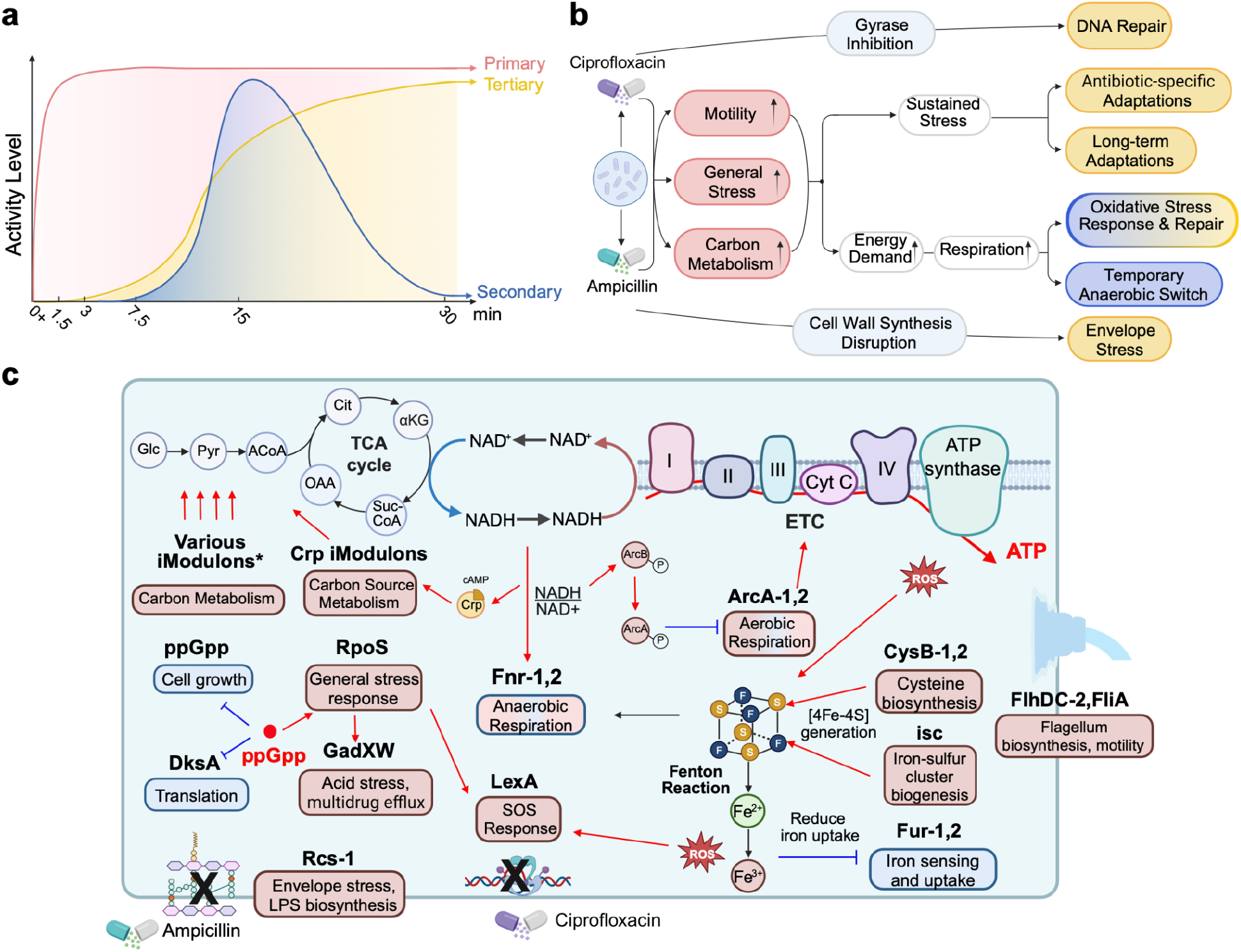
E. coli’s early response to antibiotic exposure. (Created with Biorender.com)^17^ **a)**. Diagram that outlines the timing and magnitude of the primary, secondary, and tertiary responses of E. coli in the first 30 minutes of antibiotic exposure. The 0+ time point represents samples taken immediately after the antibiotic was added (See Methods). **b)**. Proposed cascade of cellular events following antibiotic exposure. The colors of the box describing the regulatory responses are consistent with the colors representing primary, secondary, and tertiary responses in panel a. **c)**. Major responses in the first 30 minutes of antibiotic exposure outlined by key iModulons. The text in the rounded box indicates the cellular function of the associated iModulon. The color of the box displays the relative activity of the iModulon, with red representing overall activation and blue representing overall repression. The phases in which these iModulons are involved can be found in panel b and Supplementary Figure 2. The “Various iModulons” include the iModulons associated with carbon metabolism that are in Figure 3.

The primary response occurs almost immediately after antibiotic exposure and involves genes associated with a variety of cellular functions. One of the most pronounced responses is general stress adaptation. The iModulon associated with the global stress response regulator RpoS displays significantly elevated activity. In contrast, we noted that iModulons associated with growth, including the ppGpp iModulon and the Translation iModulon, are significantly downregulated. These observations align with the “fear vs. greed” tradeoff in cellular responses, where cells prioritize stress response over growth under distress (Supplementary Figure 1)^18^. These three iModulons associated with the fear-greed tradeoff contributed substantially to the gene expression variations in the antibiotic exposure data, accounting for 15.5% of the observed variance in gene expression.

In addition to activating global stress pathways, we observe increased activity in motility-associated iModulons. While resistant *E. coli* strains typically downregulate motility-associated structures to conserve energy^19–21^, the early response appears to favor “flight” over “fight”. The high activity of flagella-related iModulons suggests increased motility, while iModulons associated with biofilm formation and attachment remain repressed. This prioritization of motility over sessile adaptations represents an initial survival strategy, enabling the cells to evade antibiotic stress before committing to long-term resistance mechanisms. iModulon analysis also reveals rapid changes in the expression of genes associated with cellular structures, metal homeostasis, and notably, metabolism. The activity of all the iModulons for the antibiotic treated samples can be found in Supplementary Figure 2, and discussions on other interesting iModulons from the primary response can also be found in Supplementary Note 2. Given the high energetic demands of the early defenses, we observe widespread metabolic shifts accompanying these responses, emerging as another fundamental aspect of the primary response. These metabolic adaptations will be explored next before we turn to the secondary and tertiary phases, where the transcriptional landscape evolves and becomes more specialized and adaptive. The major transcriptional changes outlined by key iModulons are presented in Figure 2c and will be discussed in detail in the following sections.

### iModulons Uncover Detailed Metabolic Remodeling

Increased energy demands and a need to maintain metabolic flexibility under stress prompt *E. coli* to undergo extensive metabolic shifts as a primary response to antibiotic exposure. Carbon metabolism and nucleotide biosynthesis are significantly reprogrammed, accompanied by an increase in aerobic respiration and energy production.

The cells appear to be programming their metabolism to access available nutrients, as evidenced by increased activity in iModulons related to alternative carbon source utilization and anaplerosis. These iModulons are enriched in carbon source transporters and pathways that feed different carbon into glycolysis and the TCA cycle. Figure 3a maps these iModulons onto carbon metabolic pathways and illustrates how they facilitate the incorporation of different carbon sources into central carbon metabolism. In contrast, iModulons associated with certain carbon sources are downregulated (Figure 3b). This includes β-glucosides, which wild-type *E. coli* cannot metabolize, as well as other carbon sources that may be deprioritized due to differences in utilization preferences^22^. The activation of alternative carbon utilization demonstrates *E. coli*’s metabolic flexibility, ensuring a steady supply of energy and metabolic intermediates by accessing multiple carbon sources. Several iModulons associated with the cyclic AMP receptor protein (CRP) further contribute to this adaptability by linking central carbon metabolism to processes such as fatty acid and aromatic compound degradation, as well as nitrogen and phosphate metabolism. This metabolic flexibility enables *E. coli* to sustain metabolic flux despite stress-induced disruptions while efficiently adjusting to changing nutrient availability.

**Figure 3.**
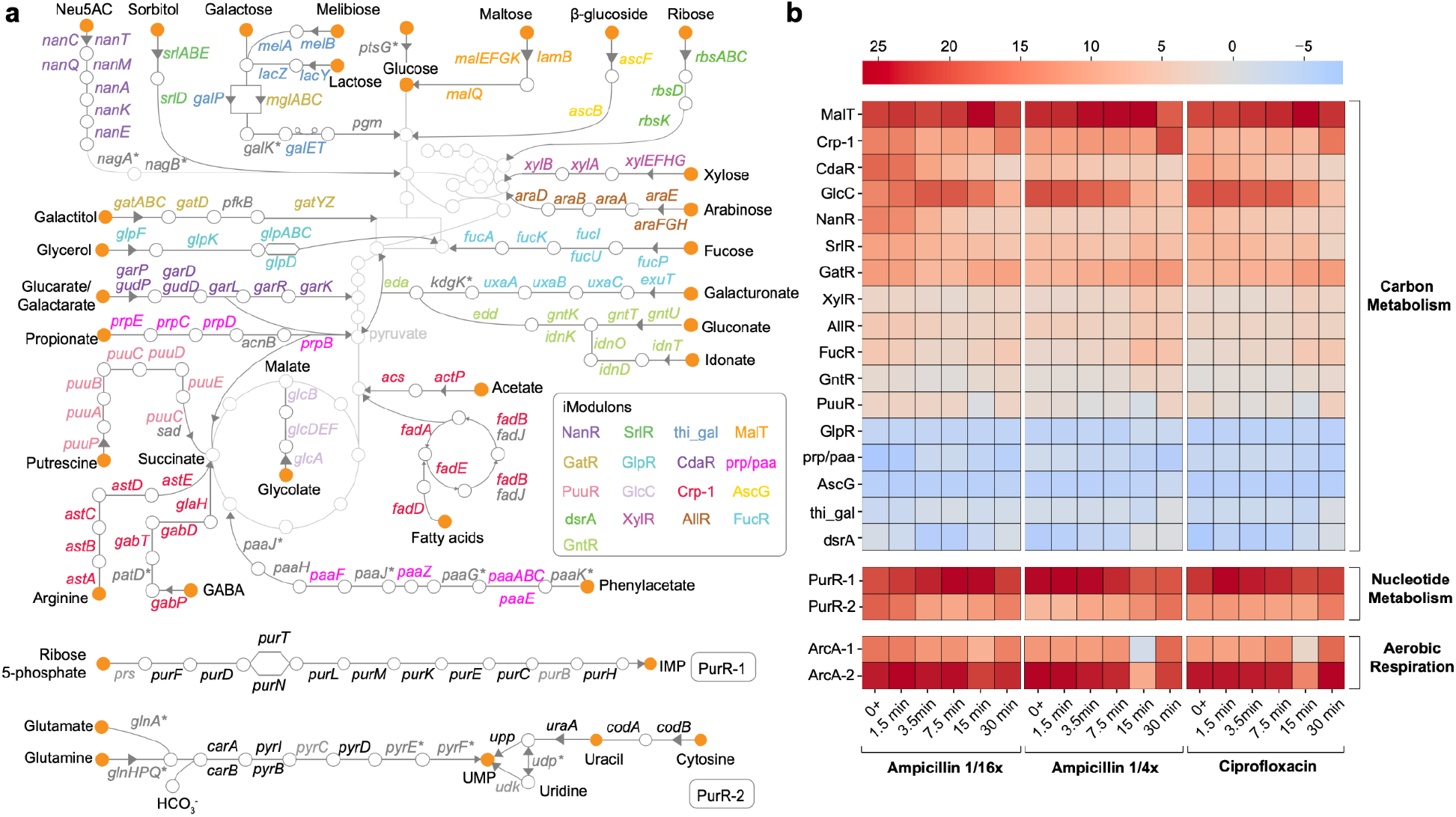
Metabolic Remodeling Revealed by iModulon Activity Changes. **a)**. iModulons with significant activity changes upon antibiotic exposure mapped onto E. coli’s metabolic network^23^. Gene names are color-coded to distinguish membership in different iModulons. Among these iModulons, the prp/paa iModulon contains both positively and negatively weighted genes mapped onto the metabolic network (the prp genes are positively weighted in the iModulon and the paa genes are negatively weighted), while the thi_gal iModulon includes only negatively weighted genes mapped onto the network. Their activities in panel b) should be interpreted in this context. Genes not associated with any iModulon are shown in gray, while genes belonging to iModulons not highlighted in this figure are shown in gray with an asterisk (*). **b)**. Activity level of carbon and nucleotide metabolism iModulons shown in panel a) as well as the ArcA iModulons for the antibiotics-exposed samples.

Simultaneously, iModulons associated with nucleotide metabolism and linked processes are upregulated, reflecting an increased demand for nucleotides (Figure 3a, b). This upregulation supports essential processes such as DNA repair, replication, and stress responses, as well as providing key metabolic regulators and signaling molecules. Additionally, antibiotic treatment can disrupt nucleotide pools and deplete building blocks, prompting the upregulation of nucleotide synthesis pathways in order to compensate^24,25^.

The active central metabolism fuels a surge in cellular respiration and increased energy production, supported by the high activity of the ArcA iModulons (Figure 3b). They are enriched in genes associated with the TCA cycle, oxidative phosphorylation, the electron transport chain, and other key processes in aerobic respiration (Supplementary Tables 2, 3). While these early metabolic responses allow the cell to cope with immediate stress, they also lead to the accumulation of toxic byproducts, including reactive oxygen species (ROS)^26,27^. ROS can compromise iron-sulfur clusters via the Fenton reaction, thereby producing highly reactive hydroxyl radicals that further damage cellular components. The Fur iModulons related to iron uptake show significantly reduced activities, likely as a response to increased intracellular free iron generated by the Fenton reaction (Supplementary Figure 2). Therefore, while the primary response facilitates rapid adaptation to sudden stress, it also imposes substantial burdens and is unsustainable over time. Ultimately, the cell must adopt more sustainable strategies to manage the long-term effects of stress and support survival^28,29^.

### iModulon Dynamics Reveal Transient Redox Regulation and Anaerobic Shift

The global activation of carbon utilization iModulons in the primary response enables the cell to rapidly generate energy through aerobic respiration to cope with antibiotic stress. However, this increased metabolic activity can disrupt redox balance, as redox cofactors such as NADH accumulate when glucose consumption outpaces the cell’s capacity to reoxidize reduced equivalents^30^. Specifically, the increased flux through the TCA cycle produces NADH faster than the electron transport chain can regenerate NAD+, leading to an elevated NADH/NAD+ ratio. This redox imbalance triggers redox-sensing regulators to adjust gene expression and restore redox homeostasis^31,32^. Consistent with this, several iModulons governed by key regulators that are sensitive to redox signals—including ArcA, Fnr, and Crp—exhibit sharp activity changes by 15 minutes post-exposure. We hypothesize that this transient shift, which stabilizes later for some iModulons, constitutes a “redox reset” to counteract the redox disturbances induced by the primary response (Figure 4a).

**Figure 4.**
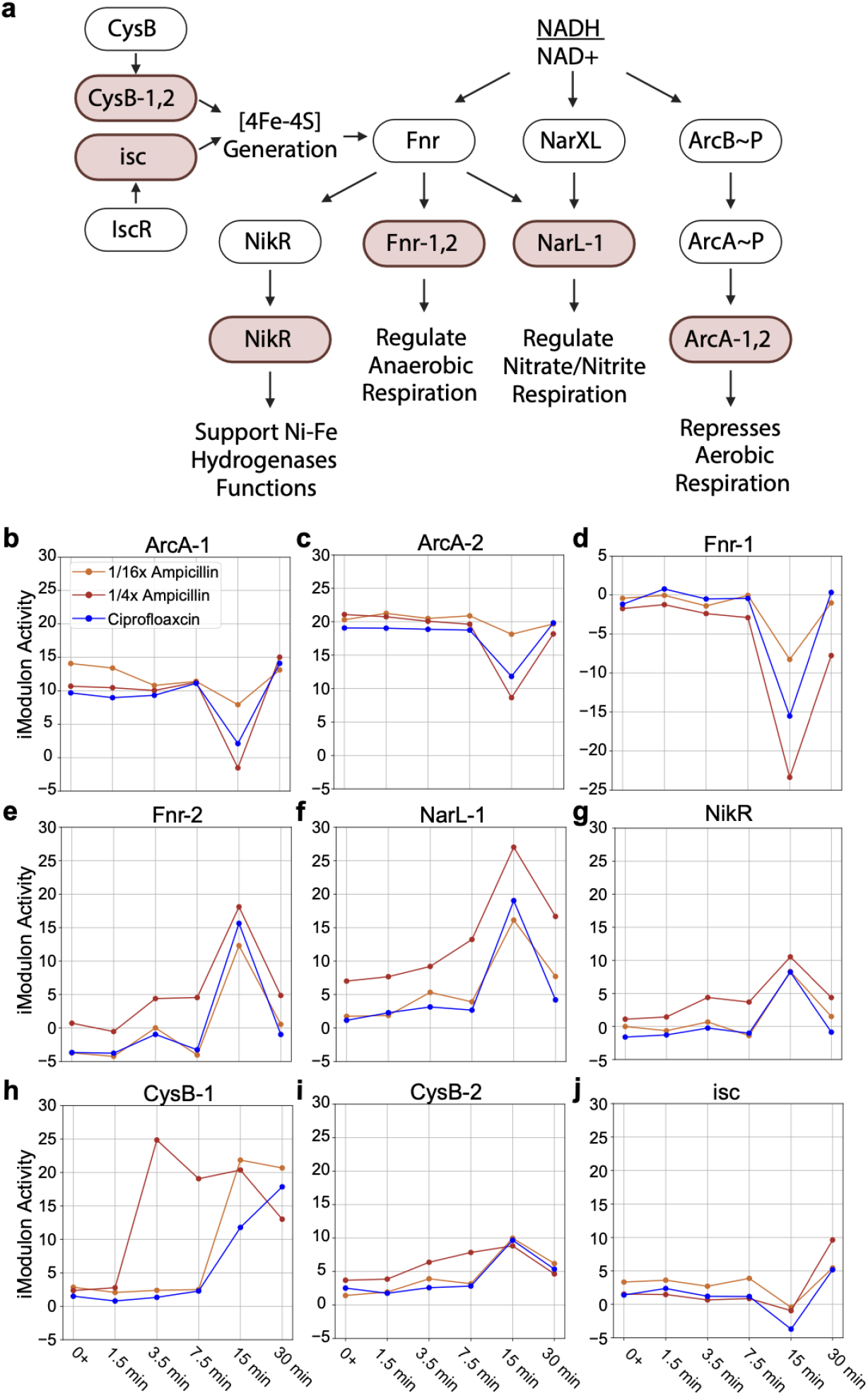
Key iModulons display temporal activity patterns. **a)**. Mechanistic outline of the secondary response with the associated regulators and iModulons. The colored boxes represent iModulons, and the black and white boxes represent regulators (created with Biorender.com^17^). **b) – j)**. iModulons associated with secondary response show significant activity changes over time. Note that genes in the Fnr-2 iModulons are negatively weighted, therefore an increase in iModulon activity reflects a “more significant decrease” in gene expression^13^.

A high NADH/NAD+ ratio triggers the phosphorylation of the ArcAB system, leading to ArcA-mediated repression of aerobic respiration genes^33^. Accordingly, we observed a sharp decrease in ArcA iModulon activities at 15 minutes (Figure 4b, c). At the same time, iModulons associated with anaerobic metabolism increased in activity, including genes regulated by Fnr and those involved in alternative electron acceptor utilization, such as DMSO and nitrite respiration controlled by NarLX (Figure 4d-f). The genes in these iModulons are shown in Supplementary Tables 4-6. This shift provides an alternative route for NADH reoxidation to NAD+, thereby restoring redox balance. The activation of Fnr also upregulates the NikR iModulon (Figure 4g), which includes the NikABCDE nickel transporter essential for NiFe hydrogenase synthesis during anaerobic growth^34,35^. We also detected increased activity in the cysteine iModulons (Figure 4h, i) and later activation of the *isc* iModulon at 30 minutes (Figure 4j), likely supporting iron-sulfur cluster biogenesis to maintain Fnr function during this phase^36,37^. While this response resembles overflow metabolism^32^, we did not observe significant changes in genes linked to acetate or lactate production. It is possible these pathways were transiently activated between 15 and 30 minutes but were not captured in our samples.

Importantly, the secondary response does not appear to be sustained, suggesting that its primary function is to correct redox imbalance rather than to serve as a long-term metabolic adaptation. Once the electron transport chain catches up or additional regulatory mechanisms adjust metabolism, the cells shift back to aerobic pathways to maximize energy efficiency. This transient activation of anaerobic pathways in response to redox imbalance has not been previously documented in *E. coli* antibiotic responses. This distinct and dynamic shift warrants further investigations to clarify the metabolic and regulatory mechanisms driving it and its role in antibiotic adaptation.

### Tertiary Response: Shift to Long-Term Adaptation

As the stress continues, the cell appears to transition from short-term corrections to longer-term adjustments supporting survival under sustained antibiotic exposure. While the primary response remains active throughout the time course and the secondary response addresses redox imbalances, we observe gradual shifts in the activity of many iModulons beginning around 7.5 minutes and persisting through 30 minutes, suggesting broad regulatory adjustments in the tertiary response. Many of the iModulons activated in the primary response evolve in this time frame. These ongoing adjustments contribute to the broader regulatory shifts that define the tertiary response and highlight how *E. coli* fine-tunes its initial defenses to meet the demands of prolonged antibiotic stress. Although these changes are distributed across a wide range of pathways (Supplementary Figure 2), one notable example is the progressive increase in Crp-related iModulons, particularly Crp-2, which is enriched in transcriptional regulators of various carbon utilization pathways (Supplementary Table 7).

Prolonged stress appears to shift metabolic regulation from broad activation toward a more selective strategy. While many alternative carbon utilization pathways remain active, certain iModulons linked to specific carbon sources—such as CdaR (glucarate/galactarate metabolism), GlcC (glycolate conversion to malate), and NanR (feeding Neu5Ac into glycolysis) — gradually decrease in activity (Supplementary Figure 2). This selective downregulation may reflect a refinement of metabolic priorities as adaptation progresses, potentially influenced by constraints such as resource allocation, metabolic efficiency, or regulatory burden. Meanwhile, Crp-2 and its associated regulators, several of which govern carbon-associated iModulons activated during the primary response, may guide this transition from global carbon activation to more coordinated, Crp-mediated metabolic remodeling. Given Crp’s role as a master regulator that integrates metabolic and redox signals, its rising activity may contribute to longer-term adjustments that balance carbon utilization with the cell’s overall redox state under sustained stress.

### iModulons Identify Antibiotic-Specific Response

The tertiary response also includes antibiotic-specific adaptations that reflect the cell’s efforts to counteract different types of antibiotic stress. The antibiotics used in this study have distinct mechanisms of action: ampicillin, a β-lactam, disrupts peptidoglycan synthesis, while ciprofloxacin, a fluoroquinolone, interferes with DNA replication by inhibiting DNA gyrase and topoisomerase IV. Consistent with these differences, we identified specific iModulons that capture unique aspects of each antibiotic’s impact on *E. coli*.

*E. coli* relies on multiple systems to manage envelope stress, each tailored to specific types of damage^38^. While many of these systems are represented in our iModulon analysis (Supplementary Figure 2), the Rcs system exhibited the most prominent signal in response to ampicillin exposure. The Rcs system (regulator of capsule synthesis) responds to disruptions in peptidoglycan biosynthesis, and plays a crucial role in resistance to β-lactams^39^. We identified two Rcs-related iModulons, with Rcs-1 exhibiting rapid and sustained activation following ampicillin exposure. Although Rcs-1 only partially overlaps with the known Rcs regulon, it includes *rcsA, rcsB*, and several genes linked to envelope stress, such as *ivy* and *mliC* (lysozyme inhibitors)^40^, *osmB* (osmotic stress resistance), *hsiJ* (novobiocin resistance), and *ygaC* (pH stress response) (Figure 5a)^41^. Notably, 14 of its 21 genes are uncharacterized y-genes, many of which are previously predicted to contribute to envelope biogenesis and have been associated with antibiotic responses^39^. We performed MIC measurements using knockout strains from the Keio collection for 17 genes in this iModulon (the remaining 4 are not represented in the collection). Most knockouts exhibited MICs comparable to the wild type or showed slightly increased susceptibility to ampicillin, while two showed modest increases in resistance (Supplementary Figure 3). Interestingly, deletion of the highest-weighted gene in this iModulon, *yaiY*, led to a notable reduction in MIC, suggesting increased vulnerability. This, combined with strong early activation of Rcs-1, supports its role in immediate defense against β-lactam-induced envelope stress (Figure 5b), making it a promising target for studying resistance mechanisms.

**Figure 5.**
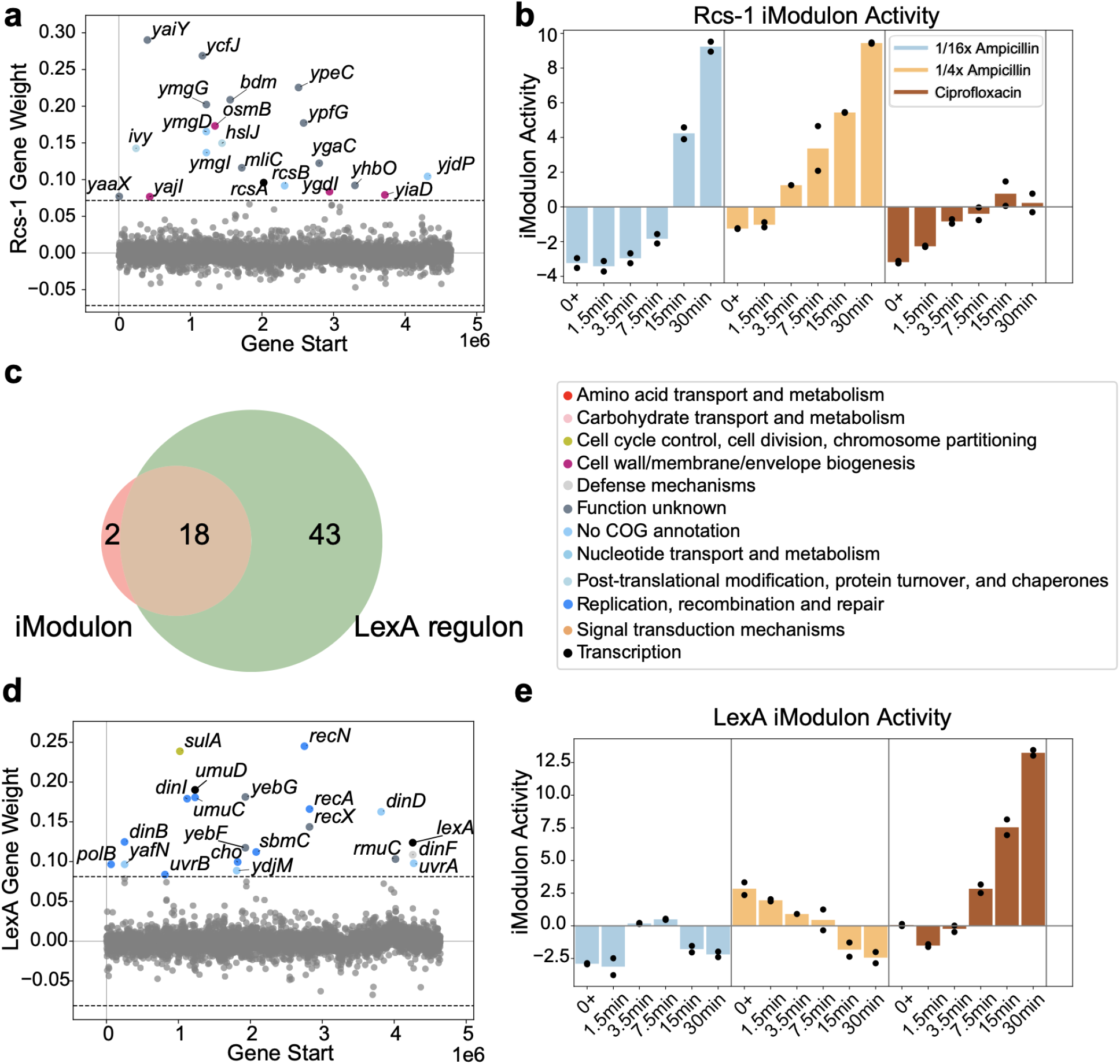
Antibiotic specific iModulon responses. **a)**. iModulon gene membership for the Rcs-1 iModulon. **b)**. The activity of the Rcs-1 iModulon for antibiotics treated samples. **c)**. Venn diagram showing the overlap between the LexA iModulon and the LexA regulon. **d)** iModulon gene membership for the LexA iModulon.**e)**. The activity of the LexA iModulon for antibiotics treated samples.

Ciprofloxacin inhibits DNA gyrase, causing DNA stress that activates the SOS response. Accumulated single-stranded DNA (ssDNA) triggers RecA, which promotes LexA self-cleavage and derepresses SOS genes^42^. In response to ciprofloxacin, we identified a LexA-associated iModulon comprising many DNA damage-inducible genes from the LexA regulon (Figure 5c, d), including *lexA, uvrA, uvrB* (nucleotide excision repair), *recA, recN* (homologous recombination), *sulA* (cell division inhibition), and the error-prone polymerases *polB, dinB*, and *umuCD* (translesion synthesis). The activity of this iModulon increases after 7.5 minutes of ciprofloxacin exposure, likely reflecting the accumulation of ssDNA beyond the activation threshold for the SOS response, and remains elevated, consistent with ongoing DNA damage from DNA gyrase inhibition. In contrast, this iModulon stays inactive during ampicillin treatment (Figure 5e), likely because β-lactams cause DNA damage indirectly through the DpiBA system, leading to a slower and less severe SOS activation^43^. Ciprofloxacin’s direct disruption of DNA rapidly induces the full SOS program captured by this iModulon, reflecting the magnitude and urgency of its DNA-targeting mechanism. Another ciprofloxacin-specific but less pronounced signal is discussed in Supplementary Note 3.

### Differential iModulon Activities Reveal Dose-Dependent Transcriptomic Responses

To assess whether transcriptional responses vary with antibiotic dosage, we compared *E. coli*’s activity under two subinhibitory concentrations of ampicillin: 1/4× MIC and 1/16× MIC. As reflected in the preceding sections, the overall regulatory and metabolic responses were largely similar, indicating that the core regulatory and metabolic responses to β-lactam stress are independent of dose at subinhibitory concentrations (Figure 1d, Figure 3b). However, higher concentrations tend to drive stronger shifts in iModulon activity, suggesting that dosage primarily affects the magnitude of adaptation rather than its regulatory structure (Figure 4b-i, Supplementary Note 4, Supplementary Figure 4). Notably, we observed selective activation of the TdcA iModulon only at the higher concentration. This iModulon contains genes in the *tdc* operon, which support threonine and serine transport and metabolism during anaerobiosis.

Similarly, the hya iModulon enriched in hydrogenase 1 genes is also activated around 15 minutes only for 1/4x MIC (Supplementary Note 4, Supplementary Figure 4). The activation of these iModulons at higher antibiotic concentrations may suggest that greater stress intensifies the cell’s metabolic demands or the anaerobic-like redox reset, leading to the engagement of additional metabolic pathways.

## Discussion

Bacterial survival under antibiotic stress relies on complex, multilayered adaptations. In this study, we applied iModulon analysis to high-resolution time-series RNA-Seq data to capture the early transcriptional response of *E. coli* to subinhibitory antibiotic exposure, revealing a structured, three-phase process of adaptation. This structure provides a genome-scale view of how regulatory programs are dynamically organized over time, offering new insight into the processes that govern early survival and adaptation under antibiotic pressure.

Our three-phase dynamic response reveals how *E. coli* manages the competing demands of antibiotic stress through a coordinated, layered strategy, with each layer addressing distinct challenges. The primary response provides a rapid, broad activation of stress responses to stabilize cellular function under sudden pressure. This global response is energetically demanding and disrupts redox balance, triggering a transient secondary response that restores homeostasis through anaerobic pathways. Meanwhile, the tertiary response establishes more sustainable adaptations, such as metabolic remodeling and antibiotic-specific defenses, to support prolonged survival as stress persists. Importantly, the responses are not strictly sequential; the sustained activity of the primary response alongside the emergence of tertiary adaptations illustrates how *E. coli* layers its defenses, maintaining immediate survival mechanisms while gradually developing more specialized, long-term strategies. Some iModulons contribute to multiple phases, with their activities dynamically shifting to support this transition and reflect the developing demands of ongoing antibiotic stress.

Beyond defining this general framework, our analysis uncovers regulatory features that merit further investigation. The strong, ampicillin-specific activation of the Rcs-1 iModulon points to an envelope stress response enriched with largely uncharacterized *y*-genes, suggesting potential roles in β-lactam tolerance that remain unexplored. The identification of previously unannotated genes in a key antibiotic response pathway provides an opportunity to refine functional annotations and potentially identify biomarkers for emerging resistance. Additionally, the selective activation of the *tdc* operon and *hya* genes at higher ampicillin concentrations—one of the few dose-dependent differences observed—may indicate an additional layer of metabolic adaptation under more severe or prolonged stress, potentially linked to sustained anaerobic-like conditions.

Together, these findings offer a new perspective on the early stages of bacterial adaptation to antibiotics. By dissecting transcriptional complexity into a structured, three-phase process, this work provides a unifying view of how *E. coli* coordinates different regulatory programs to balance immediate survival with sustained adaptation. Beyond defining this progression, our results also highlight the underlying biological processes that drive each phase, shedding light on how metabolic, redox, and stress-response systems interact under antibiotic pressure. This may help guide combination therapies by identifying vulnerabilities in adaptive stress responses that could be targeted alongside antibiotics to enhance efficacy and prevent persistence. Future studies can assess whether this framework is a general feature of bacterial stress responses or varies across different classes of antibiotics, bacterial strains—including clinical isolates—and longer timeframes. Additionally, integrating transcriptomic data with metabolomics and pangenomics could provide further insight into metabolic shifts and the genetic determinants that shape regulatory variability across *E. coli* lineages.

## Methods

### Culture Conditions and Sample Preparation

An overnight *E. coli* MG1655 culture was inoculated from a glycerol stock and grown at 37°C with shaking. The next morning, a fresh culture in MHB-CA was prepared at an initial OD_600_ of 0.05. Once the OD_600_ reached 0.5, cultures were aliquoted into plastic tubes (16 mL per tube) containing stir bars and placed on a 24-well magnetic heat block. Ampicillin (final concentration: 16 µg/mL or 4 µg/mL) or ciprofloxacin (0.016 µg/mL) was immediately added. Samples were collected in duplicate at 0, 1.5, 3.5, 7.5, 15, and 30 minutes post-exposure, treated with RNAprotect Bacteria Reagent, and stored at -80°C for RNA-Seq library preparation.

### RNA Extraction and Library Preparation

Total RNA was extracted using a Luna Nanotech PuroMAG™ Total RNA Purification Kit (NKM051-384) which included a DNaseI treatment and a modification to incorporate bead beating as a means of cell lysis. RNA quality and concentration were assessed using an Agilent TapeStation and a Nanodrop, respectively. One microgram of total RNA underwent ribosomal RNA removal using RiboRid^44^, and the resulting rRNA-depleted RNA was used to make RNA-Seq libraries with the Kapa Biosystems (Roche) RNA HyperPrep Kit, following the manufacturer’s protocol.

### Data Preprocessing

Quality control of the antibiotic exposure dataset was performed following step_3 of the iModulonMiner workflow^14^. One sample was filtered out during quality control (ampicillin 1/4x MIC 7.5 minutes) due to low number of mapped reads and poor replicate correlation. To improve the quality of the ICA decomposition with more data, we combined our antibiotic-treated MG1655 samples with MG1655 expression data from the PRECISE 1K compendium^15^.

Specifically, we filtered the PRECISE 1K dataset to retain only samples from strain MG1655, resulting in 578 samples that were merged with our dataset prior to normalization. All samples were normalized to a project reference. Because the 0-minute time points in the antibiotic exposure dataset were collected immediately after antibiotic addition (0+), and transcriptional changes during processing time might be captured, we used control samples from antibiotic-free CAMHB cultures from the study done by Sastry et al. in the PRECISE 1K dataset as the reference for this project^45^. These samples were generated using the same strain, condition, protocol, and personnel, helping to minimize technical variability between datasets. Previous analyses have shown that samples collected under these standardized conditions display high consistency across studies^15,16^. Further discussion of normalization choices, control sample consistency, and batch effects is provided in Supplementary Note 5.

### Extracting and Characterizing iModulons

ICA calculations and iModulon extractions were performed following the iModulonMiner workflow^14^. The optimal dimensionality of the dataset was determined to be 200, resulting in 132 iModulons^46^. iModulon enrichments against known regulons from RegulonDB and metabolic pathways from KEGG as well as explained variance calculations were conducted using the PyModulon Python package. iModulons are named with the associated regulator or inspected individually and named based on their gene content and functional enrichment results.

## Data Availability

The RNA-Seq data generated in this study have been deposited to the Gene Expression Omnibus (GEO) with the accession number GSE295494. All code and data used to generate the results in this paper can be found on GitHub (https://github.com/AnnieYuan21/antibiotICA-MG1655). The general iModulon analysis pipeline can be found at https://github.com/SBRG/iModulonMiner^14^.

## Acknowledgements

This work was funded by the Y.C. Fung Endowed Chair in Bioengineering at UC San Diego to B.O.P. We acknowledge seed funding in support of this work from the Tata Institute for Genetics and Society (TIGS) Endowment Fund.

We thank Dr. Donghui Choe for insightful discussions of the results and helpful experimental advice. We also thank Dr. Daniel Zielinski for valuable feedback and constructive discussion.

## References

1. Salam, Md. A. et al. Antimicrobial Resistance: A Growing Serious Threat for Global Public Health. Healthcare 11, 1946 (2023).

2. Kohanski, M. A., Dwyer, D. J. & Collins, J. J. How antibiotics kill bacteria: from targets to networks. Nat. Rev. Microbiol. 8, 423–435 (2010).

3. Deter, H. S., Hossain, T. & Butzin, N. C. Antibiotic tolerance is associated with a broad and complex transcriptional response in E. coli. Sci. Rep. 11, 6112 (2021).

4. Dwyer, D. J. et al. Antibiotics induce redox-related physiological alterations as part of their lethality. Proc. Natl. Acad. Sci. 111, E2100–E2109 (2014).

5. Martínez, J. L. & Rojo, F. Metabolic regulation of antibiotic resistance. FEMS Microbiol. Rev.35, 768–789 (2011).

6. Rychel, K., Sastry, A. V. & Palsson, B. O. Machine learning uncovers independently regulated modules in the Bacillus subtilis transcriptome. Nat. Commun. 11, 6338 (2020).

7. Poudel, S. et al. Revealing 29 sets of independently modulated genes in Staphylococcus aureus, their regulators, and role in key physiological response. Proc. Natl. Acad. Sci. 117, 17228–17239 (2020).

8. Chauhan, S. M. et al. Machine Learning Uncovers a Data-Driven Transcriptional Regulatory Network for the Crenarchaeal Thermoacidophile Sulfolobus acidocaldarius. Front. Microbiol. 12, (2021).

9. Yoo, R. et al. Machine Learning of All Mycobacterium tuberculosis H37Rv RNA-seq Data Reveals a Structured Interplay between Metabolism, Stress Response, and Infection. mSphere 7, e00033–22 (2022).

10. Yuan, Y. et al. Pan-Genome Analysis of Transcriptional Regulation in Six Salmonella enterica Serovar Typhimurium Strains Reveals Their Different Regulatory Structures. mSystems 7, e00467–22 (2022).

11. Shin, J., Rychel, K. & Palsson, B. O. Systems biology of competency in Vibrio natriegens is revealed by applying novel data analytics to the transcriptome. Cell Rep. 42, 112619 (2023).

12. Yuan, Y. et al. Machine learning reveals the transcriptional regulatory network and circadian dynamics of Synechococcus elongatus PCC 7942. Proc. Natl. Acad. Sci. 121, e2410492121 (2024).

13. Sastry, A. V. et al. The Escherichia coli transcriptome mostly consists of independently regulated modules. Nat. Commun. 10, 5536 (2019).

14. Sastry, A. V. et al. iModulonMiner and PyModulon: Software for unsupervised mining of gene expression compendia. PLOS Comput. Biol. 20, e1012546 (2024).

15. Lamoureux, C. R. et al. A multi-scale expression and regulation knowledge base for Escherichia coli. Nucleic Acids Res. 51, 10176–10193 (2023).

16. Catoiu, E. A. et al. iModulonDB 2.0: dynamic tools to facilitate knowledge-mining and user-enabled analyses of curated transcriptomic datasets. Nucleic Acids Res. gkae1009 (2024) doi:10.1093/nar/gkae1009.

17. Scientific Image and Illustration Software | BioRender. https://www.biorender.com/.

18. Dalldorf, C. et al. The hallmarks of a tradeoff in transcriptomes that balances stress and growth functions. mSystems 9, e00305–24.

19. Rundell, E. A., Commodore, N., Goodman, A. L. & Kazmierczak, B. I. A Screen for Antibiotic Resistance Determinants Reveals a Fitness Cost of the Flagellum in Pseudomonas aeruginosa. J. Bacteriol. 202, e00682–19 (2020).

20. Sharma, D., Garg, A., Kumar, M., Rashid, F. & Khan, A. U. Down-Regulation of Flagellar, Fimbriae, and Pili Proteins in Carbapenem-Resistant Klebsiella pneumoniae (NDM-4) Clinical Isolates: A Novel Linkage to Drug Resistance. Front. Microbiol. 10, (2019).

21. Lyu, Z., Yang, A., Villanueva, P., Singh, A. & Ling, J. Heterogeneous Flagellar Expression in Single Salmonella Cells Promotes Diversity in Antibiotic Tolerance. mBio 12, 10.1128/mbio.02374-21 (2021).

22. Harwani, D. Regulation of gene expression: Cryptic β-glucoside (bgl) operon of Escherichia coli as a paradigm. Braz. J. Microbiol. 45, 1139–1144 (2015).

23. Kanehisa, M., Furumichi, M., Sato, Y., Ishiguro-Watanabe, M. & Tanabe, M. KEGG: integrating viruses and cellular organisms. Nucleic Acids Res. 49, D545–D551 (2021).

24. Lopatkin, A. J. & Yang, J. H. Digital Insights Into Nucleotide Metabolism and Antibiotic Treatment Failure. Front. Digit. Health 3, 583468 (2021).

25. Yang, J. H. et al. A white-box machine learning approach for revealing antibiotic mechanisms of action. Cell 177, 1649-1661.e9 (2019).

26. Belenky, P. et al. Bactericidal antibiotics induce toxic metabolic perturbations that lead to cellular damage. Cell Rep. 13, 968–980 (2015).

27. Stokes, J. M., Lopatkin, A. J., Lobritz, M. A. & Collins, J. J. Bacterial Metabolism and Antibiotic Efficacy. Cell Metab. 30, 251–259 (2019).

28. Händel, N., Schuurmans, J. M., Brul, S. & ter Kuile, B. H. Compensation of the Metabolic Costs of Antibiotic Resistance by Physiological Adaptation in Escherichia coli. Antimicrob. Agents Chemother. 57, 3752–3762 (2013).

29. Zampieri, M. et al. Metabolic constraints on the evolution of antibiotic resistance. Mol. Syst. Biol. 13, 917 (2017).

30. van Hoek, M. J. & Merks, R. M. Redox balance is key to explaining full vs. partial switching to low-yield metabolism. BMC Syst. Biol. 6, 22 (2012).

31. Holm, A. K. et al. Metabolic and Transcriptional Response to Cofactor Perturbations in Escherichia coli. J. Biol. Chem. 285, 17498–17506 (2010).

32. Vemuri, G. N., Altman, E., Sangurdekar, D. P., Khodursky, A. B. & Eiteman, M. A. Overflow Metabolism in Escherichia coli during Steady-State Growth: Transcriptional Regulation and Effect of the Redox Ratio. Appl. Environ. Microbiol. 72, 3653–3661 (2006).

33. Park, D. M., Akhtar, Md. S., Ansari, A. Z., Landick, R. & Kiley, P. J. The Bacterial Response Regulator ArcA Uses a Diverse Binding Site Architecture to Regulate Carbon Oxidation Globally. PLoS Genet. 9, e1003839 (2013).

34. Myers, K. S. et al. Genome-scale Analysis of Escherichia coli FNR Reveals Complex Features of Transcription Factor Binding. PLoS Genet. 9, e1003565 (2013).

35. Rowe, J. L., Starnes, G. L. & Chivers, P. T. Complex Transcriptional Control Links NikABCDE-Dependent Nickel Transport with Hydrogenase Expression in Escherichia coli. J. Bacteriol. 187, 6317–6323 (2005).

36. Fontecave, M., Py, B., Ollagnier de Choudens, S. & Barras, F. From Iron and Cysteine to Iron-Sulfur Clusters: the Biogenesis Protein Machineries. EcoSal Plus 3, 10.1128/ecosalplus.3.6.3.14 (2008).

37. Mettert, E. L. & Kiley, P. J. Coordinate Regulation of the Suf and Isc Fe-S Cluster Biogenesis Pathways by IscR Is Essential for Viability of Escherichia coli. J. Bacteriol. 196, 4315–4323 (2014).

38. Mitchell, A. M. & Silhavy, T. J. Envelope stress responses: balancing damage repair and toxicity. Nat. Rev. Microbiol. 17, 417–428 (2019).

39. Laubacher, M. E. & Ades, S. E. The Rcs Phosphorelay Is a Cell Envelope Stress Response Activated by Peptidoglycan Stress and Contributes to Intrinsic Antibiotic Resistance. J. Bacteriol. 190, 2065–2074 (2008).

40. Liang, Y., Zhao, Y., Kwan, J. M. C., Wang, Y. & Qiao, Y. Escherichia coli has robust regulatory mechanisms against elevated peptidoglycan cleavage by lytic transglycosylases. J. Biol. Chem. 299, 104615 (2023).

41. Huja, S. et al. Fur Is the Master Regulator of the Extraintestinal Pathogenic Escherichia coli Response to Serum. mBio 5, 10.1128/mbio.01460-14 (2014).

42. Little, J. W. & Mount, D. W. The SOS regulatory system of Escherichia coli. Cell 29, 11–22 (1982).

43. Miller, C. et al. SOS Response Induction by ß-Lactams and Bacterial Defense Against Antibiotic Lethality. Science 305, 1629–1631 (2004).

44. Choe, D. et al. RiboRid: A low cost, advanced, and ultra-efficient method to remove ribosomal RNA for bacterial transcriptomics. PLOS Genet. 17, e1009821 (2021).

45. Sastry, A. V. et al. Machine Learning of Bacterial Transcriptomes Reveals Responses Underlying Differential Antibiotic Susceptibility. mSphere 6, 10.1128/msphere.00443-21 (2021).

46. McConn, J. L., Lamoureux, C. R., Poudel, S., Palsson, B. O. & Sastry, A. V. Optimal dimensionality selection for independent component analysis of transcriptomic data. BMC Bioinformatics 22, 584 (2021).

