## Supplementary Materials for "Revealing Transcriptomic Responses in *Escherichia coli* During Early Antibiotic Exposure"

#### Supplementary Notes

##### Supplementary Note 1 - Clarification on iModulon Artifacts

Since our dataset combines the PRECISE 1K and antibiotic exposure time series data, we identified a small number of iModulons with high variance that appear to result from technical differences between datasets rather than meaningful biological activity. The X matrix in PRECISE 1K contains 4257 genes while the antibiotic exposure series has 4305 genes. iModulons Unchar_2 consists exclusively of the gene differences between these two datasets. It does not reflect a biological signal and was therefore excluded from downstream analysis to ensure that our interpretations focus on genuine regulatory signals. In addition, we removed the csp iModulons whose activity patterns are likely influenced by specific samples in the dataset and do not reflect consistent or biologically interpretable responses (also see Methods section and Supplementary Note 5). By filtering out these artifacts, we aimed to maintain a robust and reliable analysis of the transcriptional adaptations to antibiotic stress.

##### Supplementary Note 2 - Other Notable iModulons in Primary Response

Temperature-stress related iModulons exhibited notable activity patterns in response to antibiotic exposure. The RpoH and RpoE iModulons, associated with heat shock and exocytoplasmic stress respectively, showed increased activities. Previous studies have shown that both RpoH and RpoE influence *E. coli*'s response to antibiotics[1,2], and connections between antibiotic response and temperature have been explored[3–5]. Interestingly, while ampicillin and ciprofloxacin are shown to have distinct temperature-related properties in long-term exposure, our iModulon analysis of short-term exposure revealed a heat shock-like response for both antibiotics. We hypothesize that in early exposure, both antibiotics, despite their distinct mechanisms of action, induce cellular stresses such as envelope stress and protein misfolding directly or indirectly. These challenges pose similar threats to those encountered during heat shock, as opposed to cold shock, which primarily affects membrane fluidity and protein synthesis. In this context, heat shock proteins offer broad protective effects against various types of cellular damage not limited to heat-induced stress, which makes them useful for combating diverse stresses induced by antibiotics. Concurrently, the activation of membrane-related iModulons suggests that cells are likely attempting to modify their membrane and LPS composition, potentially to reduce antibiotic flux and enhance cellular impermeability, further contributing to their defense against antibiotic stress.

Our analysis also identified two Fur iModulons and the ryhB iModulon, related to enterobactin synthesis and siderophore transport regulated by Fur, which were significantly downregulated in antibiotic-treated samples. This pattern likely reflects a cellular strategy to limit free iron and reduce oxidative damage caused by the Fenton reaction under antibiotic stress. Fur, the master regulator of iron homeostasis in *Escherichia coli*, has been extensively linked to antibiotic resistance[6,7], with prior studies showing that Fur inactivation in *E. coli* BW25113 promotes the evolution of resistance[8]. These findings reinforce Fur's central role in coordinating stress responses during antibiotic exposure, with regulatory influence that extends beyond iron metabolism.

One notable example of this broader regulatory impact is the YmfT iModulon, which was activated in antibiotic-treated samples and contains genes from the cryptic e14 prophage. This prophage has been shown to support *E. coli* survival under adverse conditions, including acid stress, and contributes to traits such as biofilm formation. Its increased activity under antibiotic stress is consistent with previous reports that cryptic prophages, including e14, enhance *E. coli* resistance to quinolones and β-lactam antibiotics.

Interestingly, the *fur* gene itself is present within the YmfT iModulon but with an opposing weighting relative to the prophage genes (Supplementary Figure 5). This suggests a potential negative regulatory role for Fur on these prophage genes. This suggests a potential negative regulatory relationship between Fur and prophage expression. Although this specific interaction has not been described in *E. coli*, iron availability is known to influence prophage activation in other bacterial species[9,10]. Given Fur's central role in managing iron homeostasis and its integration with pathways controlling oxidative stress, energy metabolism, and virulence, these observations suggest that Fur may also play a role in modulating cryptic prophage activity as part of the coordinated cellular response to antibiotic stress[11].

#### Supplementary Note 3 - Ciprofloxacin-Specific Responses Related to Histidine

For ciprofloxacin-exposed samples, we observed a subtle yet specific reduction in the activity of the histidine biosynthesis iModulon, which contains genes from the *his* operon (Supplementary Figure 6). This reduction is intriguing given the known relationship between DNA supercoiling and *his* operon expression. Ciprofloxacin's inhibition of DNA gyrase leads to decreased negative supercoiling, which can potentially derepress the *his* operon, increasing histidine biosynthesis[12,13]. However, it also causes significant DNA damage, such as replication fork stalling. This damage can create physical barriers to the transcription machinery, particularly in supercoiling-sensitive regions like the *his* operon[14]. Furthermore, maintaining a certain level of chromosomal superhelicity is crucial for proper *his* operon regulation, and disruption of this balance may contribute to the observed deactivation of the histidine iModulon[15]. These factors collectively might explain the observed low iModulon activity seen in ciprofloxacin-treated samples. As ciprofloxacin exposure persists, the cell activates various SOS responses and DNA repair mechanisms. We previously described the LexA-associated iModulon becoming active after about 7.5 minutes, representing one of these repair responses. This cellular adaptation is reflected in the gradual increase in the histidine iModulon's activity after 15 minutes, suggesting a partial restoration of his operon transcription as DNA repair processes take effect and the cell adjusts to the altered DNA topology.

### Supplementary Note 4 - Dose-dependent Transcriptional Responses

To further investigate the impact of antibiotic concentration on transcriptional responses, we compared iModulon activities between the two tested ampicillin concentrations at matched time points (Supplementary Figure 4). These comparisons show that the overall patterns of iModulon activity are highly similar between concentrations, supporting the idea that *E. coli* relies on a consistent set of regulatory and metabolic programs to respond to β-lactam stress, regardless of dose within the subinhibitory range. However, the degree of activation and repression differs between concentrations. Several stress-related iModulons, such as RpoS, as well as those involved in envelope stress and DNA repair, exhibit stronger activation at the higher concentration. Similarly, iModulons including ppGpp and Translation, which reflect the decrease in growth associated with the fear/greed tradeoff, show deeper repression at the higher dose. Additionally, the Fur-2 iModulon is consistently more repressed at the higher concentration, likely reflecting increased oxidative stress and enhanced Fenton reactions, which require tighter control of iron uptake to limit further damage.

Differences between concentrations are evident during the secondary response. While both concentrations trigger the redox reset, the lower concentration displays a less pronounced decrease in ArcA activity and weaker activation of anaerobic pathways (Supplementary Figure 4e). In contrast, the higher concentration engages a more comprehensive anaerobic program during this phase, as reflected by the stronger activation of TdcA and hya, two iModulons associated with anaerobic metabolism. The TdcA iModulon includes genes from the *tdc* operon, which support threonine and serine metabolism under anaerobic conditions, while the hya iModulon contains genes from the *hya* operon, which synthesizes uptake hydrogenase isoenzyme 1, a key component of anaerobic hydrogen metabolism.

These differences are likely driven by the greater cellular damage imposed at higher ampicillin concentrations, which intensifies stress on essential processes such as envelope integrity, DNA stability, and redox balance. In response, *E. coli* amplifies the activity of shared regulatory programs, increasing the strength of stress mitigation and metabolic adjustments without engaging fundamentally different pathways within this subinhibitory range. This pattern shows how *E. coli* can flexibly adjust the strength of its existing defenses to match the severity of antibiotic stress, maintaining an effective response without needing to rewire its underlying regulatory strategy.

### Supplementary Note 5 - Data Normalization and Control Sample Evaluation

In the antibiotic exposure dataset, the 0 minute time points were collected immediately after antibiotic addition (0+). Including handling and processing time, the samples may already capture the earliest transcriptional changes. To provide an antibiotic-free baseline for normalization, we used CAMHB no-antibiotic control samples from the PRECISE 1K dataset (Sastry et al.)[16], which were generated under the same experimental conditions, by the same researcher, and using the same experimental protocol. The PRECISE 1K study has previously demonstrated that control samples produced under these conditions show highly consistent gene expression profiles and iModulon activities[17], supporting their use as a reliable reference.

Recent analyses on the updated iModulonDB platform show that for the samples from Sastry's study, most iModulons exhibit consistent activity across samples from different projects conducted under the same protocol[18]. The csp iModulon shows higher activity levels out of consistency, and for this reason, it was excluded from downstream analysis in this study.

We additionally generated an alternative iModulon structure using the average of the 0+ samples from the antibiotic exposure dataset as the normalization reference. This alternative approach produced highly consistent iModulon structures (overall Pearson R correlation > 90%) and activity trends, with strong agreement particularly in the secondary and tertiary phases of the response. However, the 0+ samples may already capture immediate gene expression shifts. When using them as the reference, these early signals are effectively subtracted out during normalization, reducing the ability to detect the full extent of primary response dynamics. For this reason, we selected the PRECISE 1K CAMHB controls as the reference for our primary analysis. The alternative iModulon structure is available in our GitHub repository for reference.

### Supplementary Figures

#### Supplementary Figure 1

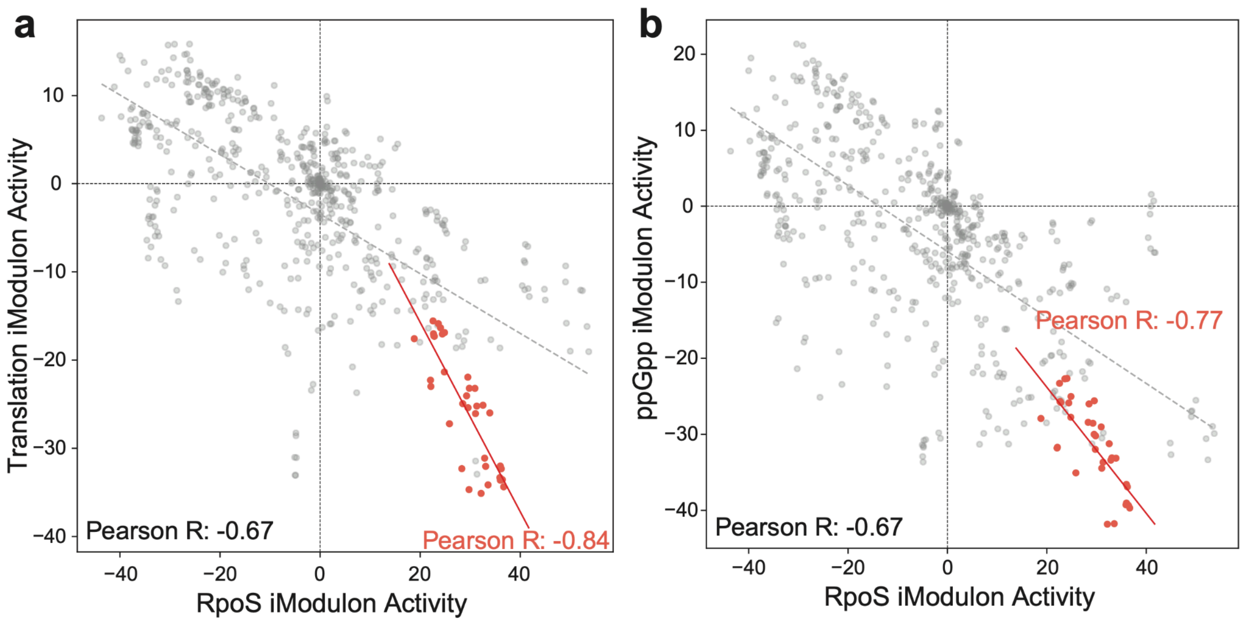


***Supplementary Figure 1. The fear-greed tradeoff. a).*** *iModulon activity comparisons of the RpoS and the Translation iModulon.* ***b).*** *iModulon activity comparisons of the RpoS and the ppGpp iModulon. The samples in the antibiotics exposure series dataset are colored in red.*

### Supplementary Figure 2

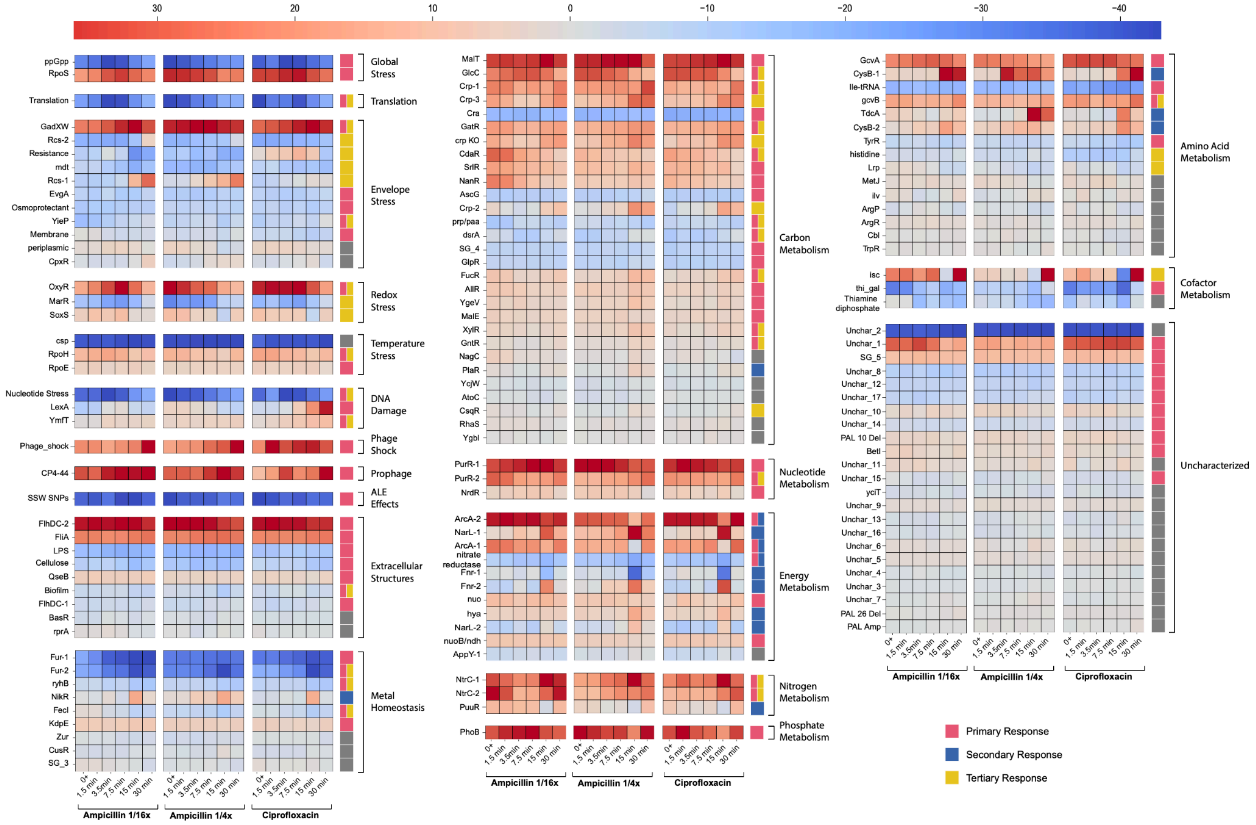


***Supplementary Figure 2. The activities of all iModulons for the antibiotic-exposed samples.*** *iModulons are grouped by functional categories. For each functional category, iModulons are sorted by high to low explained variance in the antibiotic exposure study. The color of the rightmost bar indicates the phases the iModulons are involved in. iModulons involved in multiple phases are indicated by multiple colors.*

### Supplementary Figure 3

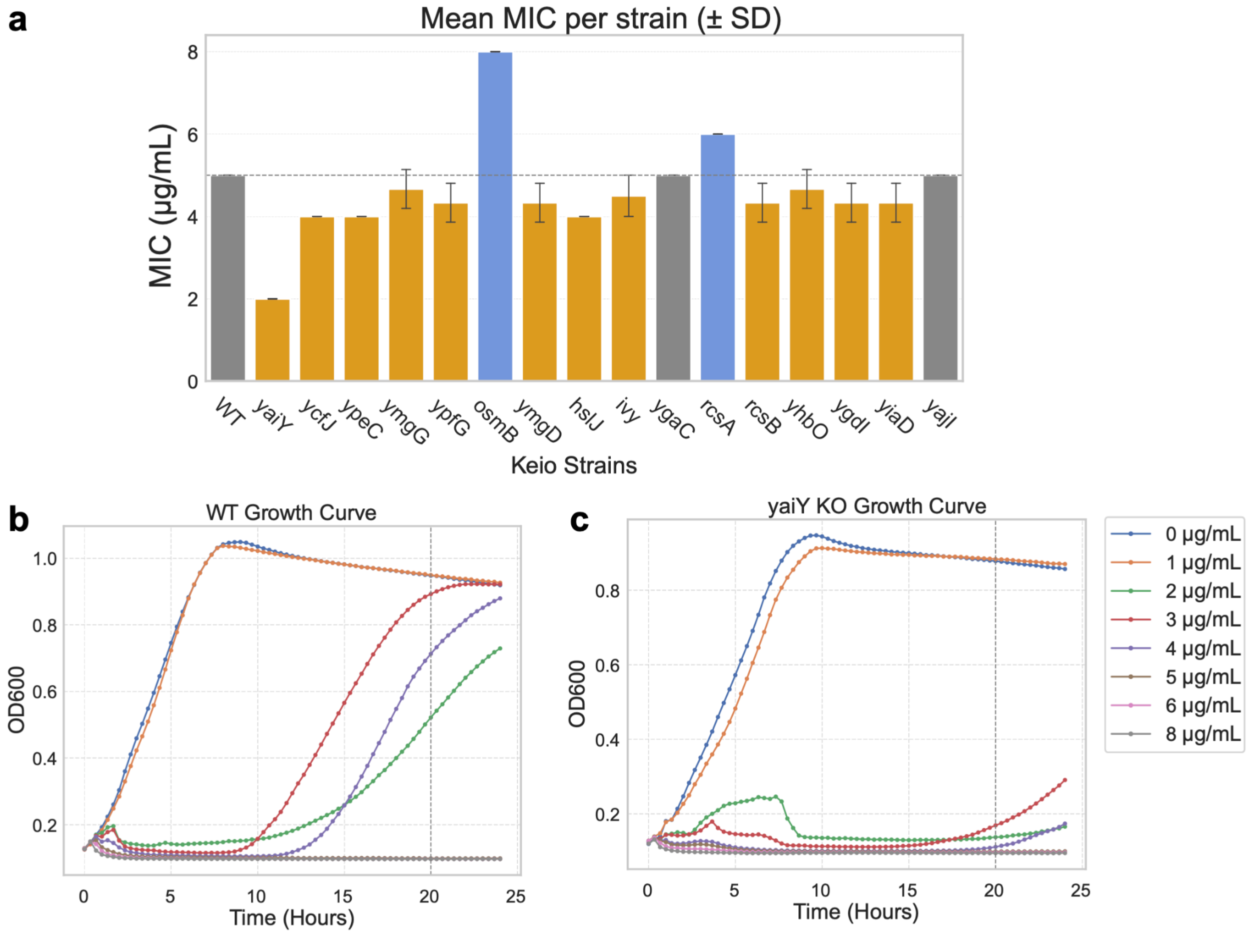


***Supplementary Figure 3. Measured MICs for knockout strains of genes in the Rcs-1 iModulon from the Keio collection. a).*** *Average MICs for each strain measured across different replicates. Color gray indicates the same MIC value as the wildtype strain. Orange represents a lowered MIC value (increase in susceptibility) and blue represents an elevated MIC value (increase in resistance). The bars for the knockout strains from left to right are arranged in the order of decreasing gene weights in the iModulon.* ***b).*** *Growth curves of the wildtype BW25113 strain under the exposure of different concentrations of ampicillin. The MIC value is determined at 20 hours (represented by a vertical dashed line).* ***c).*** *Growth curves of the* yaiY *knockout strain under the exposure of different concentrations of ampicillin.*

### Supplementary Figure 4

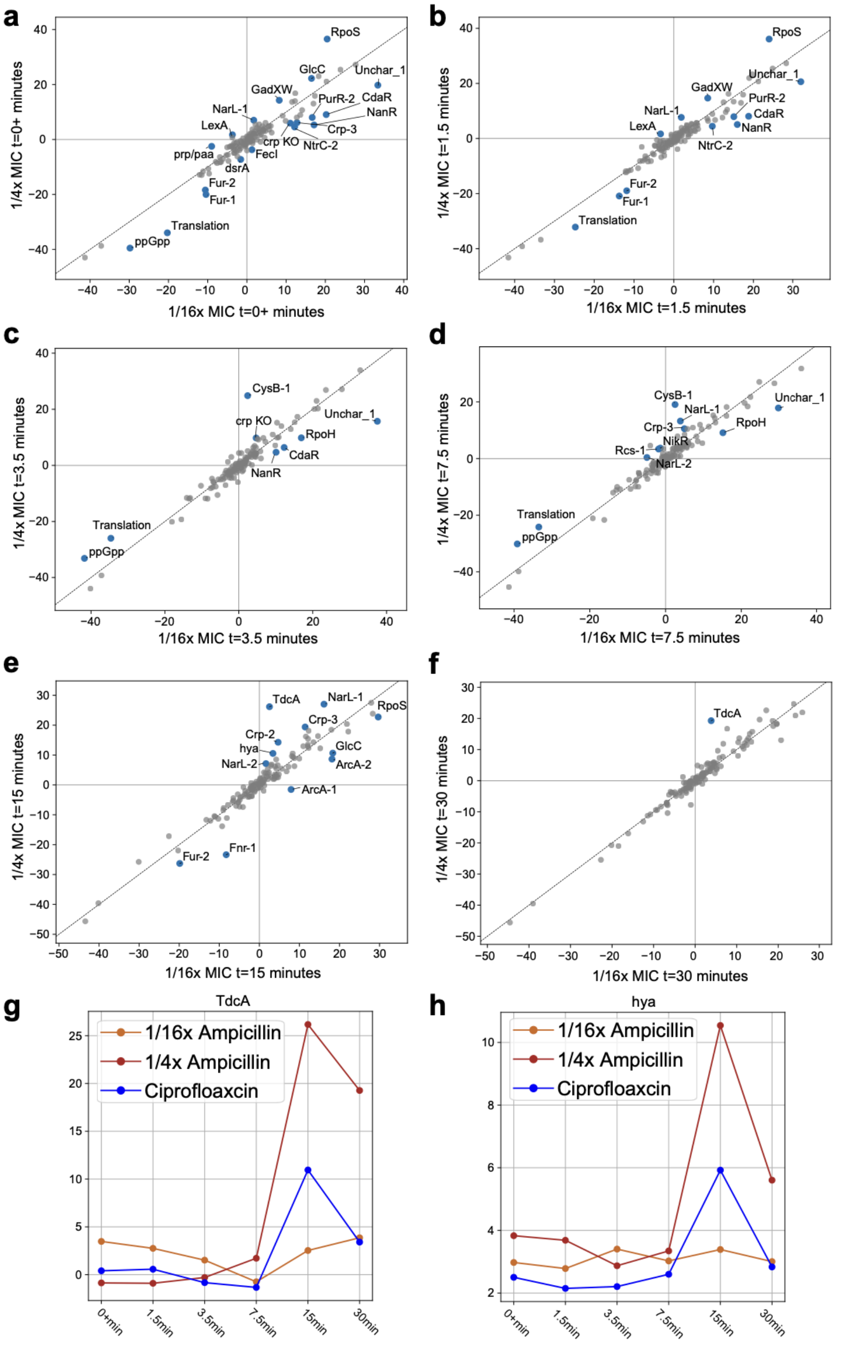


***Supplementary Figure 4. Differential iModulon activities between two concentrations of ampicillin. a-f).*** *Differential iModulon Activity (DIMA) plots for two ampicillin concentrations at each time point in the time series. The x axis shows iModulon activity at 1/16x MIC, and the y axis shows iModulon activity at 1/4x MIC. iModulons with activity differences greater than five between the two concentrations are colored in blue.* ***g).*** *Temporal activity pattern of the TdcA iModulon.* ***h).*** *Temporal activity pattern of the hya iModulon.*

Supplementary Figure 5

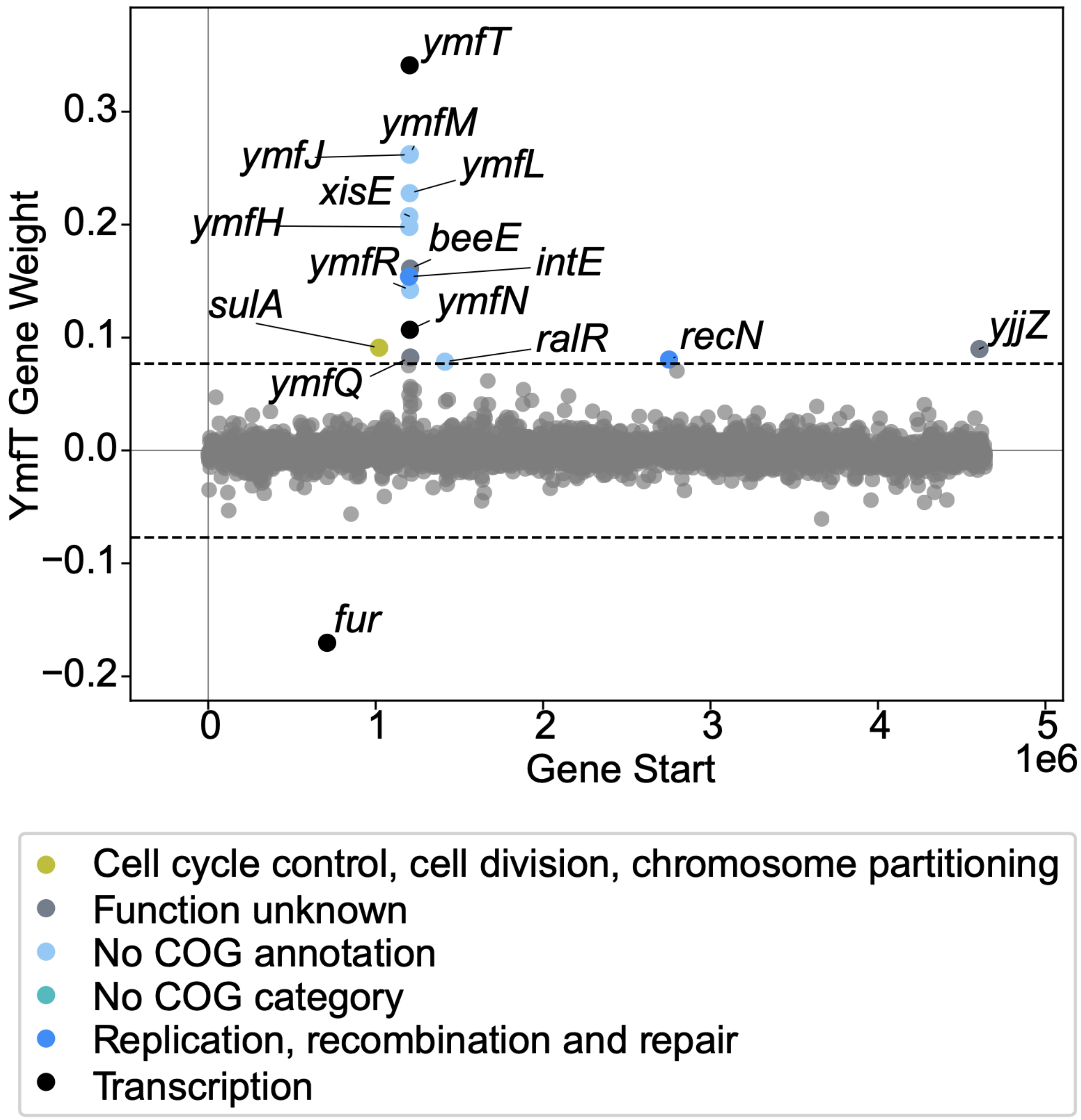


**Supplementary Figure 5. iModulon Membership and Gene Weight Plot of the YmfT iModulon**

Supplementary Figure 6

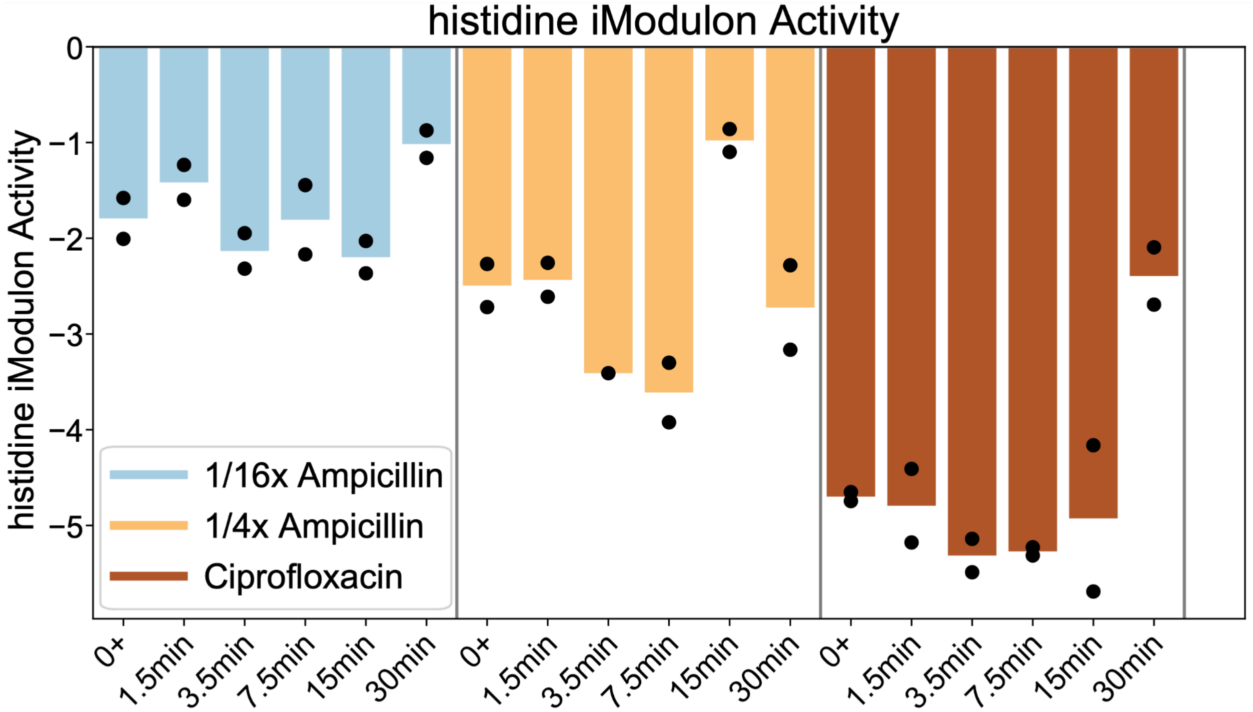


**Supplementary Figure 6. Activity of the histidine iModulon**

### Supplementary Tables

#### Supplementary Table 1 - iModulon Table

#### Supplementary Table 2 - ArcA-1 iModulon Gene Composition

#### Supplementary Table 3 - ArcA-2 iModulon Gene Composition

#### Supplementary Table 4 - Fnr-1 iModulon Gene Composition

#### Supplementary Table 5 - Fnr-2 iModulon Gene Composition

#### Supplementary Table 6 - NarL-1 iModulon Gene Composition

#### Supplementary Table 7 - Crp-2 iModulon Gene Composition

### References


1. Frizzell, J. K., Taylor, R. L. & Ryno, L. M. Constitutive Activation of RpoH and the Addition of L-arabinose Influence Antibiotic Sensitivity of PHL628 E. coli. *Antibiotics* **13**, 143 (2024).
2. Woods, E. C. & McBride, S. M. Regulation of antimicrobial resistance by extracytoplasmic function (ECF) sigma factors. *Microbes Infect.* **19**, 238–248 (2017).
3. Cruz-Loya, M. *et al.* Antibiotics Shift the Temperature Response Curve of Escherichia coli Growth. *mSystems* **6**, e00228-21.
4. Bullivant, A. *et al.* Evolution Under Thermal Stress Affects Escherichia coli's Resistance to Antibiotics. *bioRxiv* 2024.02.27.582334 (2024) doi:10.1101/2024.02.27.582334.
5. Cruz-Loya, M. *et al.* Stressor interaction networks suggest antibiotic resistance co-opted from stress responses to temperature. *ISME J.* **13**, 12–23 (2019).
6. Ezraty, B. & Barras, F. The 'liaisons dangereuses' between iron and antibiotics. *FEMS Microbiol. Rev.* **40**, 418–435 (2016).
7. Thomas, M. D. *et al.* Too much of a good thing. *Evol. Med. Public Health* **9**, 53–67 (2021).
8. Méhi, O. *et al.* Perturbation of Iron Homeostasis Promotes the Evolution of Antibiotic Resistance. *Mol. Biol. Evol.* **31**, 2793–2804 (2014).
9. Binnenkade, L., Teichmann, L. & Thormann, K. M. Iron Triggers λSo Prophage Induction and Release of Extracellular DNA in Shewanella oneidensis MR-1 Biofilms. *Appl. Environ. Microbiol.* **80**, 5304–5316 (2014).
10. Yang, G. *et al.* *Geobacter*-associated prophages confer beneficial effect on dissimilatory reduction of Fe(III) oxides. *Fundam. Res.* (2022) doi:10.1016/j.fmre.2022.10.013.

11. Seo, S. W. *et al.* Deciphering Fur transcriptional regulatory network highlights its complex role beyond iron metabolism in Escherichia coli. *Nat. Commun.* **5**, 4910 (2014).
12. Rudd, K. E. & Menzel, R. his operons of Escherichia coli and Salmonella typhimurium are regulated by DNA supercoiling. *Proc. Natl. Acad. Sci. U. S. A.* **84**, 517–521 (1987).
13. Winkler, M. E. & Ramos-Montañez, S. Biosynthesis of Histidine. *EcoSal Plus* **3**, 10.1128/ecosalplus.3.6.1.9 (2009).
14. Drlica, K. & Zhao, X. DNA gyrase, topoisomerase IV, and the 4-quinolones. *Microbiol. Mol. Biol. Rev.* **61**, 377–392 (1997).
15. Yao, Y. *et al.* A DnaA-dependent riboswitch for transcription attenuation of the his operon. *mLife* **2**, 126–140 (2023).
16. Sastry, A. V. *et al.* Machine Learning of Bacterial Transcriptomes Reveals Responses Underlying Differential Antibiotic Susceptibility. *mSphere* **6**, e00443-21.
17. Lamoureux, C. R. *et al.* A multi-scale expression and regulation knowledge base for Escherichia coli. *Nucleic Acids Res.* **51**, 10176–10193 (2023).
18. Catoiu, E. A. *et al.* iModulonDB 2.0: dynamic tools to facilitate knowledge-mining and user-enabled analyses of curated transcriptomic datasets. *Nucleic Acids Res.* gkae1009 (2024) doi:10.1093/nar/gkae1009.
